# Regulation of perceptual learning by chronic chemogenetic manipulation of parvalbumin-positive interneurons

**DOI:** 10.1101/2020.01.13.905257

**Authors:** J. Miguel Cisneros-Franco, Maryse E. Thomas, Itri Regragui, Conor P. Lane, Lydia Ouellet, Étienne de Villers-Sidani

**Author notes:** **Correspondence**: Etienne de Villers-Sidani, 3801 University Rm 742, Montreal, QC, H3A 2B4, J. Miguel Cisneros-Franco, 3801 University Rm 753, Montreal, QC, H3A 2B4.

## Abstract

Parvalbumin-positive (PV+) interneurons are major regulators of adult experience-dependent plasticity. Acute manipulation of PV+ cell activity before learning alters the rate of acquisition of new skills, whereas transient inactivation of PV+ cells interferes with retrieval of previously learned information. However, the effects of sustained PV+ cell manipulation throughout training remain largely unknown. Using chemogenetics in rat auditory cortex during an adaptive sound disrimination task, here we show that PV+ cells exert bidirectional control over the rate of perceptual learning. Down-regulation of PV+ cell activity accelerated learning, but increasing their activity resulted in slower learning. However, both interventions led to reduced gains in perceptual acuity by the end of training relative to controls. Furthermore, longitudinal training performance was functionally correlated with measures of neural synchrony and stimulus-specific adaptation. These findings suggest that, whereas restricting PV+ cell activity may initially facilitate training-induced plasticity, a subsequent increase in PV+ cell activity is necessary to prevent further plastic changes and consolidate learning.

## Introduction

Adult experience-dependent plasticity typically occurs in the context of goal-directed behavior, which requires sustained attention and is facilitated by reward or punishment ^1–3^. Learning-induced cortical remodeling requires a change in the local excitatory/inhibitory (E/I) balance ^4,5^, which is partially regulated by fast-spiking basket cells known as parvalbumin-positive (PV+) interneurons, the most prevalent type of inhibitory interneuron in the cortex ^6,7^.

Recent studies have demonstrated that the degree of cellular expression of the calcium-binding protein PV is a reliable proxy for cellular activity, as PV staining intensity is tightly correlated with the magnitude of experience-dependent plasticity ^8–10^. Similarly, motor and spatial memory training are associated with shifts in PV ‘network configuration’, such that initial incremental trial-and-error learning is accompanied by reduced PV expression and a low E/I synaptic-density ratio, whereas expertise is associated with increased PV expression and E/I synaptic-density ratio ^8,9^. Whether similar mechanisms regulate adult plasticity in primary sensory cortex remains unknown. And although these studies show that acute manipulation of PV+ interneuron activity can affect the rate of learning, the effects of sustained modulation of PV-inhibition on the acquisition and consolidation of new skills have not been documented.

To address these gaps in knowledge, we trained adult Long-Evans rats on an adaptive sound discrimination task and measured PV expression in primary auditory cortex (A1) at different time points during training. We then used chemogenetics to selectively and chronically manipulate PV+ cell activity in A1 during the same training task. Finally, we measured the functional outcomes of concurrent training and PV+ cell manipulation, and their relationship with longitudinal behavioral performance. Our findings shed new light on the inhibitory regulation of adult experience-dependent plasticity and have the potential to spur new lines of research aimed at improving learning in health and disease.

## Results

### Transient downregulation of PV expression during perceptual learning

In sensory systems, improved perception can be achieved through practice in specific sensory discrimination tasks, a process known as perceptual learning (PL) ^3,11^. To test whether PL is associated with changes in PV expression in A1 we trained rats on an adaptive auditory oddball discrimination task and examined the intensity of PV staining at different time points during training with post-mortem immunohistochemistry (**Figure 1A**). In this go/no-go behavioral paradigm, rats were rewarded for correctly identifying the presence of a deviant (oddball) tone in a short sequence of identical (standard) tones presented at 4 pulses per second (p.p.s.). The behavioral task followed a staircase procedure (one-up/one-down) with six levels of difficulty. Rats (Trained, n=4) were trained for 10 weeks with 5-6 one-hour sessions per week and improved steadily in performance (**Figure 1B**). Both hit rate and false-positive (FP) rate increased during early stages of training. On average, FP rate peaked at ~8 sessions, followed by a steady decline as rats learned to inhibit responses to non-target stimuli. We therefore chose this time point in training to assess the relationship between PV expression and learning during the early training phase in a separate group of rats (midT, n = 3) trained on the same task. For comparison, we also examined PV expression in a group of untrained rats (Control, n = 6).

**Figure 1.**
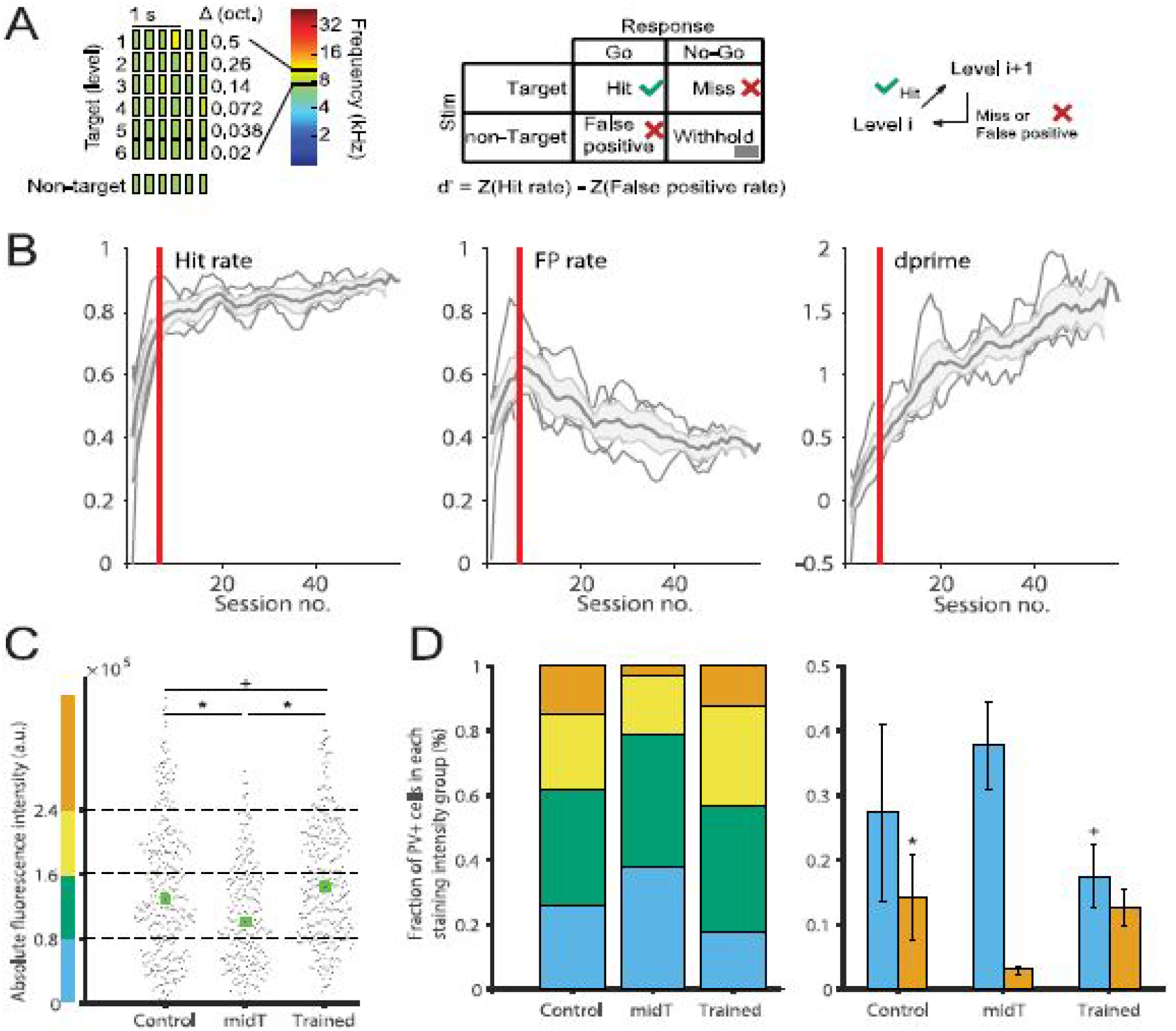
Effect of auditory discrimination training on A1 PV expression. (**A**) *Left*, task difficulty increased progressively by reducing the frequency difference (Δfreq) between oddball and standard tones with each increase in level to a maximum of 6 levels. *Right*, a correct response to an oddball-containing target stimulus (hit) led to an increase in task level, lack of response to a target (miss) or a response to the non-target stimulus (false positive) resulted in a time-out and a decrease in level (staircase one-up, one-down). (**B**) Training performance of the ‘Trained’ group. Mean hit rate, false-positive (FP) rate, and d’ over a period of 60 sessions. (**C**) Overall PV staining intensity across groups: untrained (‘Control’), mid-training (‘midT’, session no. 8, vertical lines), and ‘Trained’ (session no. 60). Effect of condition: χ2(2) = 45.01, p < 0.001, Kruskal-Wallis test; Control vs midT, p = 0, Control vs Trained, p = 0.057, midT vs Trained, p = 0, corrected with Tukey’s post-hoc test). (**D**) Cumulative (*left*) and relative (*right*) proportion of PV+ cell staining intensity for each condition. Further analysis by sub-groups of intensity levels revealed a significant effect of group on the proportion of high-intensity PV+ cells (χ2(2) = 6.54, p = 0.038; Control vs midT, p = 0.032) and a near-significant effect of group on the proportion of low-intensity PV+ cells (χ2(2) = 4.67, p = 0.09, Trained vs midT, p = 0.08). * p < 0.05, + p ≤ 0.09. Values shown are mean ± SEM. Number of subjects, measurements per group: Control, 6, 343, midT, 3, 241, Trained, 4, 324.

PV+ cells were divided into four subclasses as a function of their staining intensity ^8,12^. As previously reported in other cortical areas ^8,9^, we found downregulation of PV expression during early stages of training (**Figure 1C-D**, which likely facilitates modifications in cortical circuits that are needed for incremental trial-and-error PL ^13,14^. Conversely, expertise upon completion of training was associated with high PV expression, which might promote the consolidation of training-induced changes and prevent further circuit remodeling ^7,15^.

### Sustained PV+ cell inactivation enhances the rate of perceptual learning

Experimentally-induced changes to PV-network configuration prior to training have opposite effects such that acutely decreasing or increasing PV+ cell activity results in accelerated or delayed learning, respectively ^8^. However, the effects of sustained PV+ cell manipulation throughout training remain unknown. To tackle this question, we trained adult PV-Cre Long-Evans rats transfected with an inhibitory or excitatory DREADD [pAAV-hSYN-DIO-hM4D(Gi)-mCherry, “PV_I_” or pAAV-hSYN-DIO-hM3D(Gq)-mCherry, “PV_E_,” respectively; n = 12 each; **Supplementary Figure 1**] on the same auditory oddball discrimination task. A subset of PV_I_ and PV_E_ rats (n = 8 each) rats were treated with CNO (1 mg/kg, i.p.) 20-30 minutes before each training session, while the rest received saline injections (**Figure 2A**). Saline-treated rats did not differ in their behavioral performance and were thus analyzed together as the trained-control group (T-Ctrl, n = 8). We calculated average performance measures for training bins of 6 sessions per bin, representing approximately one week of training each.

**Figure 2.**
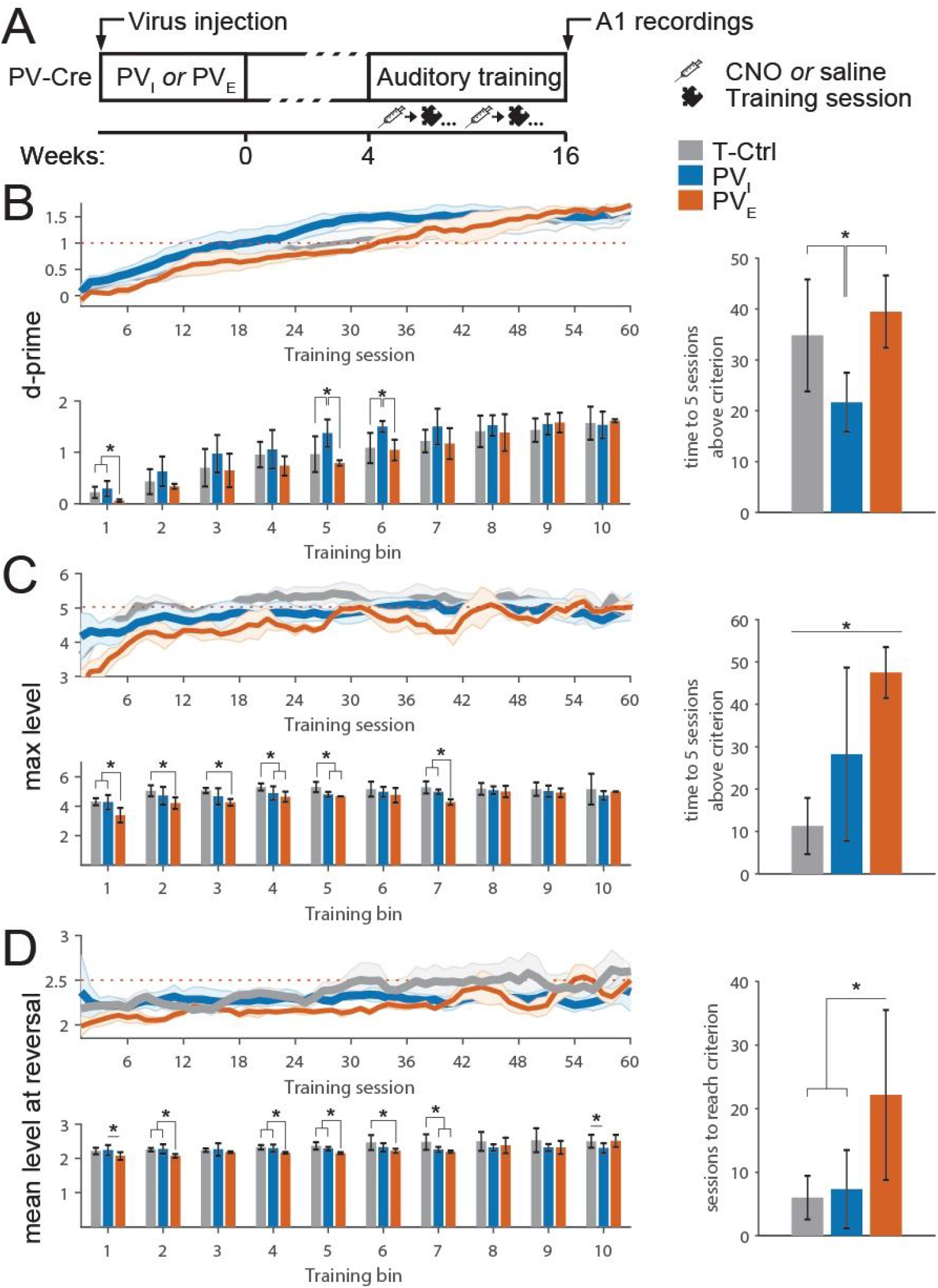
Effects of sustained A1 PV+ cell manipulation on auditory discrimination training. (**A**) *Left*, experimental timeline. *Right*, the adaptive oddball discrimination training used was the same as described in Figure 1. (**B**) *Left*, mean d’ reached over 10 training bins (6 sessions/bin) in an adaptive oddball discrimination task. The interaction was significant (F(18,135) = 2.42, p = 0.0021; repeated measures (rm) ANOVA), as were the simple main effects tests for bins 1, 5, and 6 (all p ≤ 0.03; Tukey’s post-hoc test). *Right*, number of sessions needed to reach d’ ≥ 1 (F(2,18) = 4.56, p = 0.0249, one-way ANOVA; difference relative to PV_I_ group, all p ≤ 0.037, with Tukey’s post-hoc test). (**C**) *Left*, mean maximum level reached over 10 training bins. The interaction was significant (F(18,126) = 2.49, p = 0.0172), as were the simple main effects tests for bins 1-5 and 7 (all p ≤ 0.03). *Right*, no. of sessions needed to reach level 5 or higher (F(2,19) = 48.37, p < 0.001, all pairwise comparisons p ≤ 0.002). (**D**) *Left*, mean level at reversal over 10 training bins. The interaction was significant (F(18,126) = 3.37, p = 0.002), as were the simple main effects tests for bins 1, 2, 4-7, and 10 (all p ≤ 0.04). *Right*, number of sessions needed to reach mean level at reversal ≥ 2.5 (F(2,20) = 7.64, p = 0.004, PV_E_ vs T-Ctrl/PV_I_, both p ≤ 0.016). * p < 0.05

We first calculated the sensitivity index d’ to estimate detection of the target stimuli during each session, where d’ ≥ 1 indicates successful detection. d’ was significantly higher for T-Ctrl and PV_I_ rats relative to PV_E_ rats early in training (bin 1, both p ≤ 0.0289), while the PV_I_ group performed better than the rest around mid-training (bins 5 and 6, all p ≤ 0.0162; **Figure 2B**, *left*). When analyzing the number of sessions needed to reach training criterion, we found that PV_I_ rats required less sessions (21.6 ± 5.8) to reach five consecutive sessions with d’ ≥ 1 than either other group (T-Ctrl, 34.5 ± 11.2 sessions, PV_E_, 39.5 ± 6.9, both p ≤ 0.037 relative to PV_I_; **Figure 2B**, *right*).

We then analyzed performance as measured by hit rate and FP rate, the two components that determine d’ (**Supplementary Figure 2**). Differences in between-group performance around mid-training were significant only in the case of FP-rate, such that PV_I_ rats had lower FP rates than the T-Ctrl group in training bins 5 and 6 (both p ≤ 0.0367). These results suggest that during early stages of training and despite responding to the target tone at the same rate as the other groups, PV_I_ rats were better at suppressing their response to non-target sounds, which is consistent with previous reports on the role of PV+ interneurons in operant learning ^6,7^.

### Dynamic regulation of PV+ cell activity is required for optimal learning

The use of an adaptive task allowed us to compare performance at increasing levels of difficulty. The difference in the number of sessions needed to consistently reach higher difficulty levels (five consecutive sessions with maximal level ≥ 5) was significant for all between-group comparisons, with T-Ctrl rats requiring the fewest number of sessions to reach criterion (T-Ctrl: 11.2 ± 6.6, PV_I_: 28.2 ± 9.4, PV_E_: 47.5 ± 5.8; all pairwise comparisons p ≤ 0.002; **Figure 2C**, *right*). In line with these results, the T-Ctrl group reached a daily maximum level that was significantly higher than the PV_E_ group during early- and mid-training (bins 1-5 and 7; all ≤ 0.0103), and higher than the PV_I_ group around mid-training (bins 4 and 5; both p ≤ 0.0393; **Figure 2C**, *left*).

Given that our task had a one-up one-down staircase design, advancing from one level to the next was relatively easy, as it required only one correct (hit) response. Thus, the information conveyed by maximum level alone might not necessarily represent how well a subject performed during any given session. We therefore calculated the mean level at reversal points, a more sensitive measure to assess improvement in perceptual ability as it represents the difficulty level associated with ceiling in performance for any given training session ^16,17^. Again, the T-Ctrl group consistently performed better than the PV_E_ group during early- and mid-training (bins 2 and 4-7, all p ≤ 0.038), and better than the PV_I_ group late in training (bins 7 and 10, all p ≤ 0.04; **Figure 2D**, *left*). As with previous behavioral parameters, we sought to quantify the number of sessions needed to consistently reach criterion (mean level at reversal > 2.5) for each group. Because mean level at reversals oscillated between 2 and 2.5 for all groups throughout most of the training period and no rats outside the T-Ctrl group reached 5 sessions with values above 2.5, we calculated instead the number of sessions required to reach a mean level at reversals equal or greater than 2.5. Interestingly, the PV_E_ group needed three times as many sessions (22.1 ± 13.3) than the other groups to reach this criterion (T-Ctrl: 6 ± 3.4, PV_I_: 7.3 ± 6.1; both p ≤ 0.016 relative to PV_E_; **Figure 2D**, *right*).

### Prolonged PV+ cell manipulation precludes the full expression of adaptive plasticity associated with learning

Due to their extensive dendritic and axonal arborization across cortical layers ^18,19^ and their position relative to pyramidal cells, PV+ cells are capable of inhibiting neurotransmission near the site of action potential initiation ^6^. From this strategic position, PV+ interneurons contribute to recurrent and lateral inhibition ^20,21^, suggesting that they play a major role in shaping cortical receptive fields ^22^ and modulating neural synchrony ^23^. To examine the effects of concurrent training and PV+ interneuron manipulation on these functional properties, we recorded auditory evoked potentials of a dense sample of neurons covering the whole area of A1 ^24^ in naïve and trained rats

Synchronous neural activity facilitates the processing ^25^ and integration ^26^ of sensory stimuli, and has been shown to increase with PL ^27^. We measured the degree of spontaneous synchronization between A1 single units by calculating cross-correlation functions for single-unit neuron pairs recorded in silence as in ^28^ (**Supplementary Figure 3**). We found a significant effect of condition—Naïve T-Ctrl, PV_I_, PV_E_—on neural synchrony, characterized by an increase in the mean peak correlation coefficient in the T-Ctrl group, relative to naïve and PV_I_ subjects (both p ≤ 0.024; **Figure 3A,B**).

**Figure 3.**
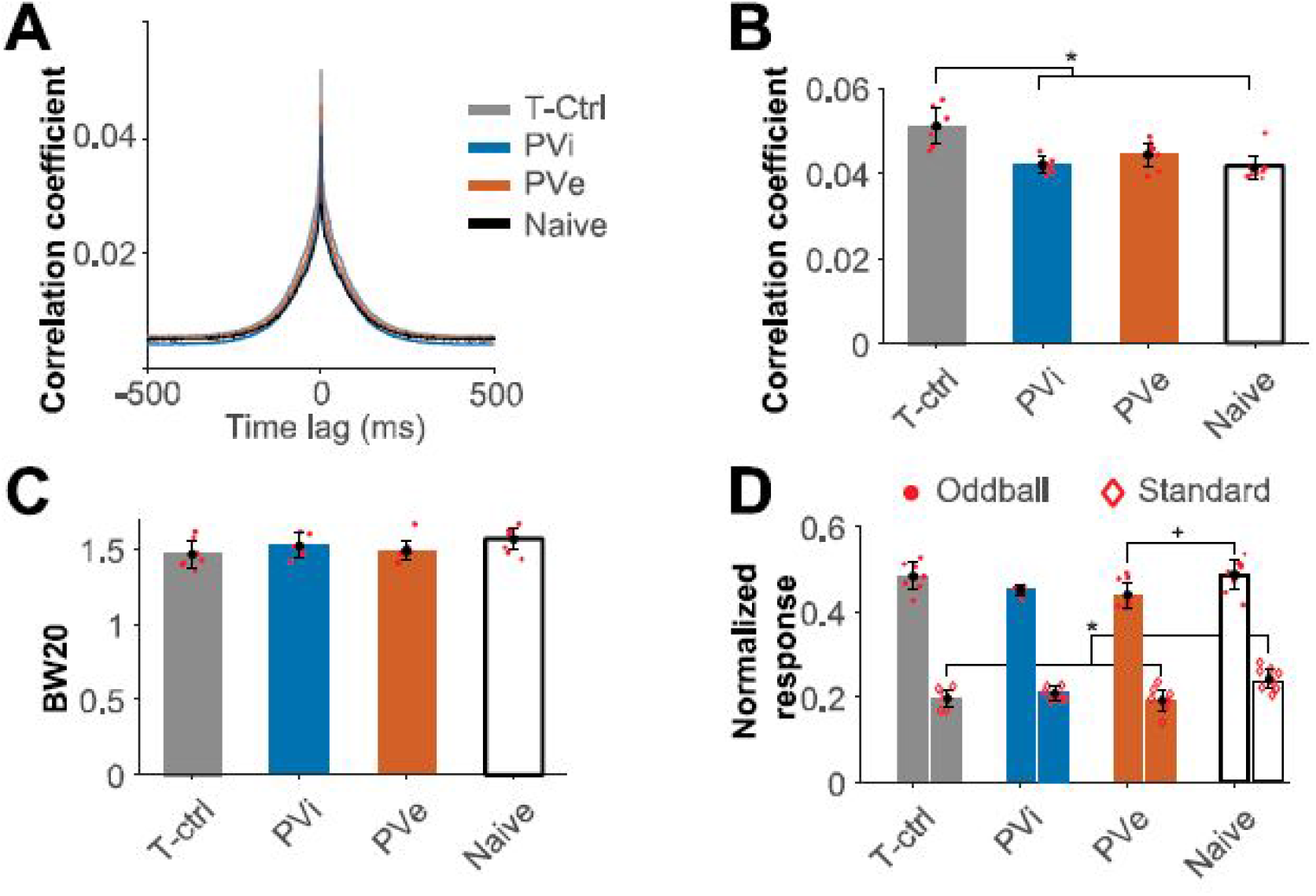
Training-related changes in cortical sensory processing. (**A**) Average cross-correlogram for all pairs < 0.5 mm apart in each experimental condition. (**B**) Peak cross-correlation coefficients (χ^2^(3) = 15.15, p = 0.0017, Kruskal-Wallis test; T-Ctrl vs naïve/PVI, both ≤ 0.024, all others, p ≥ 0.18, corrected with Tukey’s post-hoc test). (**C**) Bandwidth 20 dB SPL above auditory receptive field threshold (χ2(3) = 6.28, p = 0.09). (**D**) Stimulus-specific adaptation. Normalized responses to standard (diamonds) and oddball (circles) pure tones presented at 3 p.p.s. Standard, χ^2^(3) = 11.7, p = 0.008; naïve vs T-Ctrl/PV_E_, both p = 0.01; all others, p ≥ 0.3. Oddball, χ^2^(3) = 8.75, p = 0.032; PV_E_ vs naïve, p = 0.07, all others, p ≥ 0.11. * p <0.005

Training was not associated with any major changes in receptive field selectivity as measured by tuning bandwidth 20 dB SPL above threshold (BW20; **Figure 3C**). Although training paradigms involving the detection of a single target tone may lead to increased ^29,30^ or decreased frequency selectivity, our findings are in agreement with previous reports of normal BW values following adaptive discrimination training ^31,32^.

Neurons along the auditory pathway show a progressively reduced response to repetitive (standard) sounds, which is restored when a deviant stimulus (oddball) is presented ^33,34^. This form of deviance detection, known as stimulus-specific adaptation (SSA), is involved in the selection of sensory inputs that should be selectively suppressed during training ^35^. To examine the effect of training on SSA in A1 we used 10-min-long trains of pure tones, which included a standard (7 kHz, presented 80% of times) and five oddball tones (1.25–20 kHz, in one-octave increments). Training was associated with overall decreased responses to standard tones relative to non-trained subjects, which were significant for the T-Ctrl and PV_E_ groups (both p = 0.01; **Figure 3D**). We also found a significant effect of condition on the response to oddballs following training, but further analysis, controlling for multiple comparisons, revealed only a modest decrease in response to oddballs in the PV_E_ group, relative to naïve rats (p = 0.07).

The reduction in responses to both standard and oddball sounds in the PV_E_ group is consistent with previous research suggesting that PV+ cells contribute to SSA by providing non-specific inhibition ^36^. Taken together, these findings suggest that chronic manipulation of PV+ cell activity precluded the full expression of training-induced adaptive plasticity that is conducive to improved sensory processing ^5,37,38^. Indeed, among trained rats, only those in the T-Ctrl group exhibited both increased neural synchrony and reduced responses to standard stimuli. We therefore asked whether individual differences in training performance could account for these varied functional outcomes, as discussed in the following section.

### Learning progression and its correspondence to sensory processing

Although considerable research has been devoted to correlating electrophysiological and behavioral outcomes ^2,3,24,39^, rather less attention has been paid to the relationship between functional outcomes and longitudinal training performance ^29,30^. We considered all of d’, hit rate, FP rate, and level data obtained throughout the training period on the one hand; and BW, synchronization, and SSA measurements on the other hand. This approach, however, poses a statistical challenge, as the amount of observations (derived from dozens of training sessions and one recording session) significantly outnumbered that of experimental subjects. To address this issue, we ran a partial-least-squares multivariate analysis (PLS)—a method that has been previously used to examine the relationship between behavioral and functional data ^40^—to obtain individual behavioral (B) and functional (F) scores.

This analysis revealed a statistically significant association between B and F scores (r = 0.73; **Figure 4A**). We then investigated the contribution of individual functional parameters to overall B scores. Of the four components derived from PLS (one per functional feature), only the first component was considered, as its variance explained was greater than would be expected by chance (~38% of the covariance between electrophysiology and behavior, p = 0.05, all others, covariance ≤ 30%, p ≥ 0.37; **Supplementary Figure 4**). Whereas all four functional parameters were positively correlated with B scores, response magnitude to oddballs (**Figure 3D**) was the most strongly correlated (r = 0.70; **Figure 4B**). In other words, higher F scores (primarily driven by oddball response) were correlated with higher B scores. Note that both scores were, in general, higher for subjects in the T-Ctrl group (**Figure 4A**), which also showed a better overall performance in terms of sessions needed to reach different performance criteria (**Figure 2B-D**).

**Figure 4.**
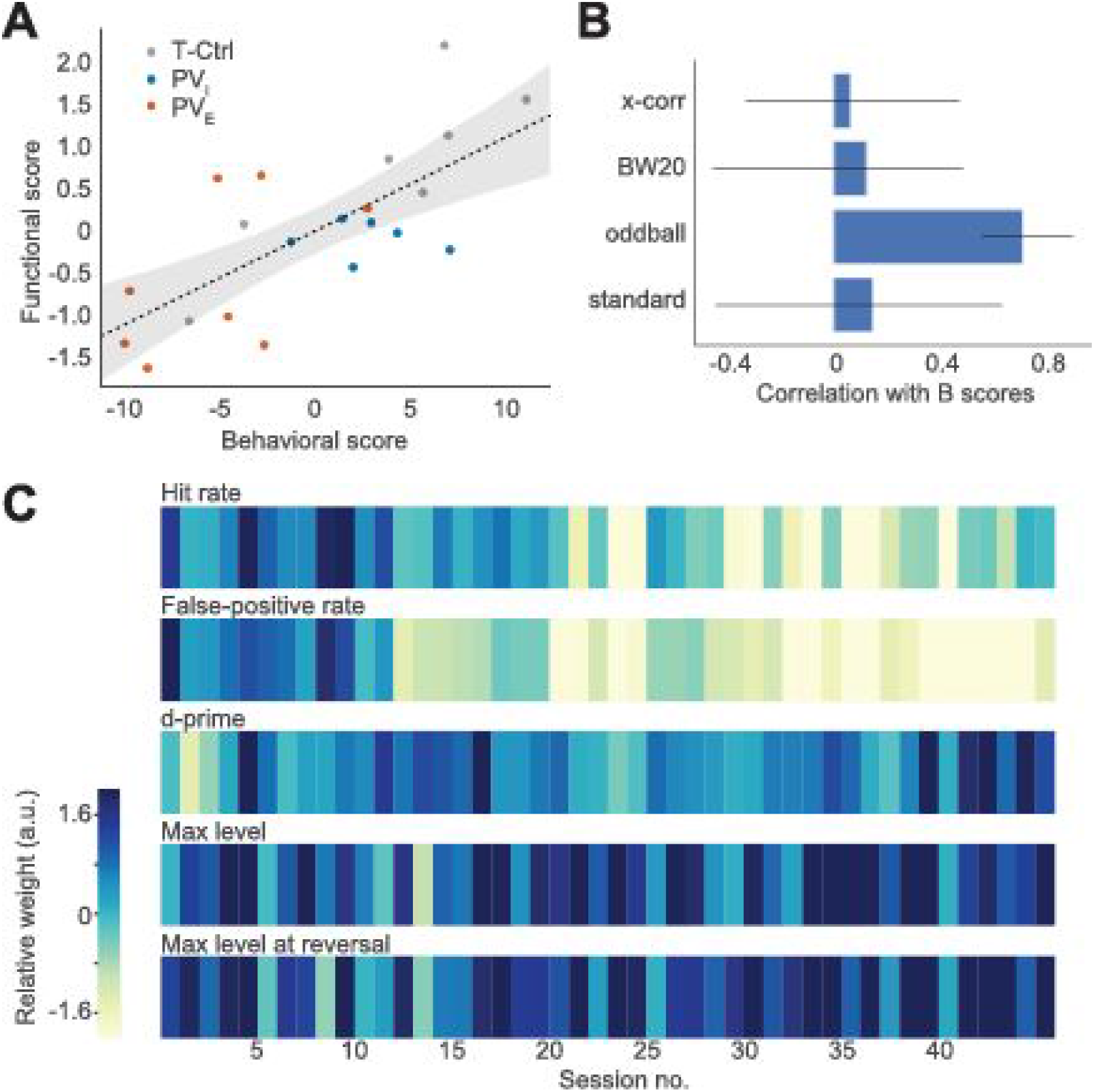
Correlation between behavioral and functional patterns. Individual behavioral and functional scores were estimated for all training sessions and one electrophysiological recording session. (**A**) Behavioral and functional scores derived from PLS analysis for each experimental subject expressed. (**B**) Correlations (i.e. loadings) of functional scores with the learning pattern. Error bars represent bootstrap-estimated 95% CIs. (**C**) Contribution of each behavioral measure to F scores. Functional connections are weighted by bootstrap ratios, a measure of reliability.

To better understand this relationship, we examined the relative contributions of each behavioral parameter to F scores. This analysis revealed that F scores tended to be positively correlated with the variables maximum level and mean level at reversal throughout the training period, and negatively correlated with FP rate, especially around mid- and late training (**Figure 4C**). These observations are in agreement with the task structure, whereby enhanced perceptual acuity (oddball detection) and lower FP rates lead to improved performance in terms of levels.

## Discussion

Our results suggest that A1 PV+ interneurons bidirectionally regulate plasticity during PL, extending previous research on other (non-sensory) learning modalities ^7,37,41^. This observation is in agreement with the biphasic model of skill learning described by Vinogradov et al. ^42^. In this model, *Phase 1* is characterized by rapid improvements in behavior, which have been associated with downregulation of cortical inhibition in animal models ^43^ and in humans ^44^. In light of previous findings that transient specific manipulation of PV+ interneurons affects the initial rate of learning ^8,37,41^, our results reinforce the mounting evidence indicating that suppression of PV+ cells is necessary for the initiation of learning ^6^.

*Phase 2*, in contrast, is characterized by modest behavioral gains coupled with functional reorganization of task-specific representations in the brain ^45,46^. In human motor cortex, for example, although this reorganization was not evident until after four weeks of training, it persisted for over 20 weeks without further training ^47^. How do these functional changes relate to PV-inhibition? Together, evidence from two independent areas of research may answer this question. First, expertise at late stages—and after the completion—of training is associated with a shift to a ‘high-PV network configuration’ ^7^, which is in agreement with reports that activation of PV+ cells improves perception ^48^. Second, although *Phase 2* is characterized by slower learning than *Phase 1* both in health and disease, this slowdown is even more pronounced in impaired neural systems, such as in autism ^49^, aging ^31^, and following noise exposure ^50^. As these conditions are also associated with altered E/I balance ^51^ and reduced PV expression ^6^, it is likely that an increase in PV+ cell function is necessary for late improvements in accuracy and for the consolidation of learning ^15,52^ (**Figure 5**).

**Figure 5.**
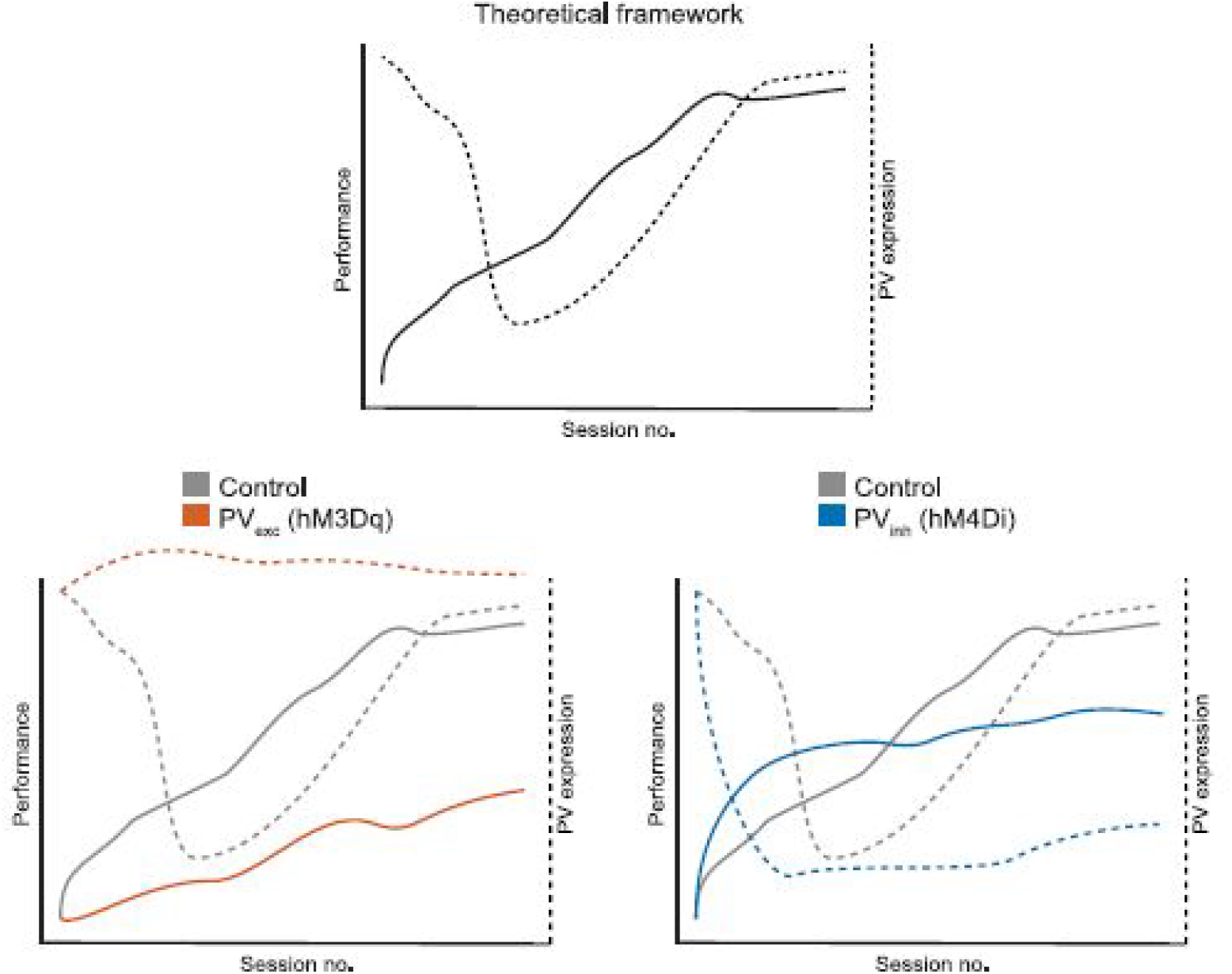
Hypothesized role of PV+ interneurons in perceptual learning. *Top*, trajectory of PV-inhibition and behavioral performance during learning. *Bottom*, hypothesized effects of chronic chemogenetic manipulation of PV+ cells.

To our knowledge, this is the first study of perceptual learning in Cre-transgenic rats. These findings thus open the door to future studies on the role of PV+ cells in learning, particularly in the context of aging, as the rat auditory system is more resilient to age-related deficits compared to mice ^53,54^. Another contribution of the present study is the use of multivariate analysis to examine the relationship between learning progression and electrophysiological measures at the individual level. Through this approach, we learned that behavioral performance and outcomes are functionally correlated with measures of neural synchrony and stimulus-specific adaptation. These findings extend previous research on the role of PV+ cells in sensory processing and demonstrate the utility of multivariate analysis in the study of adult plasticity.

Despite being innovative in terms of both the animal model and analytical approach used, certain shortcomings of our study should be addressed. First, we collected longitudinal data pertaining to behavioral performance but obtained a single readout of neuronal activity. In light of recent advances in devices for chronic and reliable neurophysiological recordings ^55^, future studies may collect both functional and behavioral data simultaneously over the whole training period to better understand the relationship between neuronal circuit function and performance. Future studies could also help determine whether these measures are correlated with PV expression. Although this matter was beyond the scope of this study, based on previous studies ^56–58^, a correspondence between the specific chemogenetic manipulation and PV staining intensity is to be expected. In addition, given that evidence to date regarding the lateralization of auditory discrimination abilities remains inconclusive, and the more general focus of the present investigation, we decided to perform bilateral virus injections. Whereas previous studies have reported impaired performance in sound localization ^59^ and discrimination of frequency-modulated tones ^60^ following lesions in either left, right, or bilateral auditory cortex, more recent investigations suggest that the processing of physiologically-relevant stimuli might indeed be lateralized ^61^. Therefore, future studies are needed to determine whether the role of PV-related inhibition on auditory discrimination abilities is lateralized.

In conclusion, these findings highlight the potential of chemogenetic approaches for harnessing cortical plasticity and open the door to future research on the inhibitory regulation of cortical plasticity during learning. These studies may also explore the effects of chronic PV+ cell manipulation on different regulatory elements of plasticity, including other interneuron subtypes and the neuromodulatory systems that control attention and motivation ^5,38,62,63^, with the ultimate goal of aiding recovery from brain injury or disease.

## Methods

### Experimental subjects

Adult PV-Cre rats were raised from commercially obtained breeding pairs (Horizon Discovery). Animals were housed on a 12-h light/12-h dark cycle and provided with ad libitum food and water unless otherwise noted. All procedures were approved by the local Animal Care Committee and follow the guidelines of the Canadian Council on Animal Care.

### Stereotaxic injections

Under isoflurane anesthesia, bilateral virus injections were performed four weeks before the beginning of chemogenetic experiments, as reported previously with Cre transgenic rats ^64^. After exposing the skull, craniotomies were made above left A1 using stereotaxic coordinates ^65^: bregma AP, −4.5mm; ML, −7mm; DV, 4.5mm. Following dura removal with a 27-G needle, 1 μL virus solution (pAAV-hSYN-DIO-hM4D(Gi)-mCherry, 5 × 10^12^ GC/mL; or pAAV-hSYN-DIO-hM3D(Gq)-mCherry, 1.2 × 10^13^ GC/mL) was injected at a depth of ~500 μm using a precision syringe pump (PHD Ultra 4400, Harvard Apparatus, Holliston, MA; injection rate, 0.05 μL/min). After injection, the needle was left in place for 10 minutes, to allow the virus to diffuse from the needle before retracting it. The surgical wound was sutured and cleaned, the rat was monitored until full recovery and medicated with antibiotic (enrofloxacin, 10 mg/kg, s.c.) and analgesic (carprofen, 5 mg/kg, s.c.) for at least 3 days ^56,57^.

### Behavioral training

The rats’ behavior was shaped in three phases. During phase A, rats were trained to make a nose poke response to obtain a food reward. During phase B, rats were trained to make a nose poke only after presentation of an auditory stimulus (single 7-kHz tone pip at 60 dB SPL). During phase C, the actual training program, rats were trained at level 1 to make a nose poke only for the target stimulus (containing an oddball frequency) and not for a foil stimulus with six identical tones of frequency identical to the standard frequency (7-kHz). The tones were presented at 60 dB SPL. Every training session started at level 1. The level was increased after one correct target identification and decreased after a response to a non-target (i.e., false-positive) or miss (one up/one-down). The task difficulty was progressively increased by reducing in exponential steps the frequency difference between standards and oddballs from 0.5 octaves (level 1) to 0.02 octaves (level 6).

Training was performed in an operant training chamber contained within a sound-attenuated chamber. Psychometric functions and stimulus target recognition thresholds were calculated for each training session by plotting the percentage of ‘go’ responses as a function of the total number of target stimuli (i.e., hit ratio) and the percentage of false positives as a function of the total number of foils (i.e., false-positive ratio). Learning curves were reconstructed by plotting maximal level reached over successive days of training.

A single behavioral trial was defined as the length of time between the onsets of two successive tone trains. The intertrial interval was selected at random from a range of 4 to 6 s. A rat’s behavioral state at any point in time was classified as either “go” or “no-go.” Rats were in the “go” state when the photobeam was interrupted. All other states were considered “no-go.” For a given trial, the rat could elicit one of five reinforcement states. The first four states are given by the combinations of responses (go or no-go) and stimulus properties (target or non-target; **Figure 1A**). Go responses within 3 s of a target were scored as a hit; a failure to respond within this time window was scored as a miss. A go response within 3 s of a non-target stimulus was scored as a false positive. The absence of a response was scored as a withhold. A hit triggered the delivery of a food pellet. A miss or false positive initiated a 5 s “time-out” period during which time no stimuli were presented. A withhold did not produce a reward or a time out.

### PV cell manipulation during training

We treated PV_I_ and PV_E_ rats with i.p. clozapine-N-oxide (CNO, 1 mg/kg) or vehicle (NaCl, 1 mL/kg) at least 20 minutes before each training session. Earlier studies of designer receptors exclusively activated by designer drugs (DREADDs) typically contrasted the effect of CNO administration between subjects that were transfected with either a DREADD or a reporter only ^66,67^. Recent findings, however, call into question the need to use CNO treatment as a negative control in chemogenetic experiments. First, CNO itself has no known biological effect and does not cross the blood-brain barrier ^68^. Indeed, the effects of CNO treatment in DREADD-transfected subjects are the result of systemic conversion to clozapine ^68^, which at low CNO doses (1 mg/kg) binds almost exclusively to DREADDs and has no known electrophysiological or behavioral effects ^69^. These reasons support the increasing use of vehicle treatment instead of CNO in control subjects (e.g., ^70,71^, the approach we chose for the present study.

### Immunohistochemistry

Immediately following the end of experiments, rats received a high dose of ketamine/xylazine/acepromazine (130/26/3 mg/kg i.p.) and were perfused intracardially with 4% paraformaldehyde in 0.1 M phosphate-buffered saline (PBS) at pH 7.2. Expression of markers of interest were examined by fluorescence immunohistochemistry using standard methods ^72^.

Immediately after perfusion, rat brains were removed and placed in the same fixative overnight for further fixation and then transferred to a 30% sucrose solution, snap-frozen, and stored at −80°C until sectioning. Fixed material was cut in the coronal plane along the tonotopic axis of A1 on a freezing microtome at 40 μm. Tissue was incubated overnight at 4°C in either monoclonal or polyclonal antisera (for anti-GABA: Sigma #A2052, dilution 1:5000; for anti-PV: Sigma #P3088, dilution 1:10 000; for mCherry: ThermoFisher #M11217, dilution 1:2000). Tissue samples were always processed in pairs during immunostaining procedures to limit variables relative to antibody penetration, incubation time, and post-sectioning age/condition of tissue. A Zeiss LSM 710 confocal microscope was used to assess fluorescence in the immunostained sections. Quantification of PV+ cell optical density was performed in Image J and MetaMorph imaging software (Molecular Devices Systems, Toronto, ON), respectively. Digital images of A1 cortical sections were taken with a 40x objective (Zeiss LSM 710). All quantification was assessed in 300-400 μm wide A1 sectors (rostral, middle, caudal) extending from layer 1 to the underlying white matter by an experimenter blind to the age of the animals. PV+ cells were classified into four subclasses according to staining intensity (a.u.) as follows: low-PV, 0-0.4 × 10^3^; intermediate low-PV, 0.4-0.8 × 10^3^; intermediate high-PV, 0.8-1.2 × 10^3^; high-PV, >1.2 × 10^3^.

### Electrophysiology

Rats were pre-medicated with dexamethasone (0.2 mg/kg) to minimize brain edema. They were anesthetized with ketamine/xylazine/acepromazine (65/13/1.5 mg/kg, i.p.) followed by a continuous delivery of isoflurane 1% in oxygen delivered via endotracheal intubation and mechanical ventilation. Vital signs were monitored using a MouseOx device (Starr Life Sciences, Holliston, MA). Body temperature was monitored with a rectal probe and maintained at 37 °C with a homeothermic blanket system. The rats were held by the orbits in a custom designed head holder leaving the ears unobstructed. The cisterna magna was drained of cerebrospinal fluid to further minimize brain edema. The left temporalis muscle was reflected, A1 was exposed and the dura was resected. The cortex was maintained under a thin layer of silicone oil to prevent desiccation.

Cortical responses were recorded with 64 channel tungsten microelectrode arrays (Neuronexus, Ann Arbor, MI). The microelectrode array was lowered orthogonally into the cortex to a depth of 470-600 μm (layers 4/5) where vigorous stimulus-driven responses were obtained. The extracellular neural action potentials were amplified, filtered (0.3-5 kHz), sorted, and monitored on-line. Acoustic stimuli were generated using TDT System III (Tucker-Davis Technologies, TDT, Alachua, FL) and delivered in a free field manner to the right ear through a calibrated speaker (TDT) located at a distance of 20-25 cm. A software package (OpenEx; TDT) was used to generate acoustic stimuli, monitor cortical response properties on-line, and store data for off-line analysis. The evoked spikes of a single neuron or a small cluster of neurons were collected at each site.

Frequency-intensity receptive fields were reconstructed by presenting pure tones of 63 frequencies (1-48 kHz; 0.1 octave increments; 25 ms duration; 5 ms ramps) at 8 sound intensities (0-70 dB SPL in 10 dB increments) in a free-field manner to the right ear at a rate of one stimulus per second. The characteristic frequency (CF) of a cortical site was defined as the frequency at the tip of the V-shaped tuning curve. For flat-peaked tuning curves, the CF was defined as the midpoint of the plateau at threshold. For tuning curves with multiple peaks, the CF was defined as the frequency at the most sensitive tip (i.e., with lowest threshold). The CF and threshold were determined using an automated routine developed in the MATLAB environment (The MathWorks Inc., Natick, MA). A1 was identified within auditory cortex based on its rostral-to-caudal tonotopy, reliable short-latency tone-evoked neuronal responses, and relatively sharp V-shaped receptive fields (Polley et al., 2007).

### Statistical analysis

Analyses were performed using Matlab 2017b. For normally distributed data, ANOVA or repeated-measures ANOVA were used. When data were not normally distributed, nonparametric tests (Kruskal-Wallis) were used. All reported p values were corrected for multiple comparisons with Tukey-Kramer test. PLS analysis was performed as described by ^40^, and the code was deposited in an online public repository ^73^. The datasets generated during and/or analysed during the current study are available from the corresponding author on reasonable request.

## Acknowledgements

This research was supported by the Canadian Institutes of Health Research (CIHR, Grant MOP-133426 to E.d.V.-S. And a CIHR Vanier Canada Graduate Scholarship to J.M.C.-F.), the Natural Sciences and Engineering Research Council of Canada (scholarship to M.E.T.), the Centre for Research on Brain, Language and Music (CRBLM, scholarship to M.E.T.), Fonds de Recherche du Québec–Nature et Technologies (FRQNT, scholarship to M.E.T. and operating grants to CRBLM), Fonds de Recherche du Québec–Société et Culture (FRQSC, operating grants to CRBLM). The authors declare no competing financial interests. We would like to thank Yohann Chaudron for his help with behavioral training, and Patrice Voss for his comments on the manuscript.

## Author contributions

**Conceptualization**: J.M.C.-F. and E.d.V.-S.; **Formal analysis**: J.M.C.-F. and I.R.; **Funding acquisition**: E.d.V.-S.; **Investigation**: J.M.C.-F., M.E.T, I.R., C.P.L., and L.O.; **Methodology**: J.M.C.-F., M.E.T., and L.O.; **Supervision**: E.d.V.-S.; **Visualization**: J.M.C.-F.; **Writing – original draft preparation**: J.M.C.-F. and I.R.; **Writing – review and editing**: J.M.C.-F., M.E.T., C.P.L., and E.d.V.-S.

## Supplementary information

**Supplementary Figure 1.**
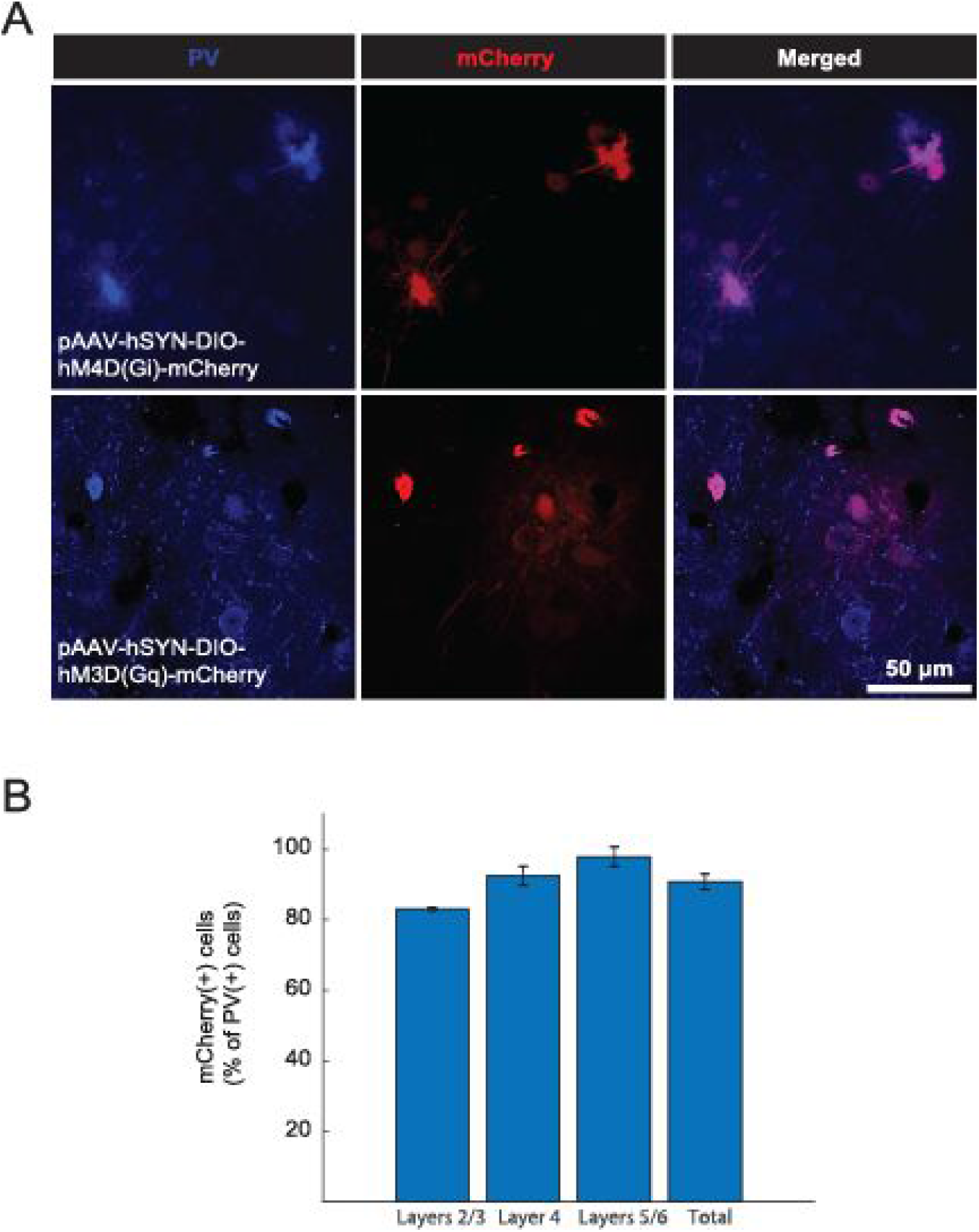
Effectiveness of DREADD transfection of PV+ cells. (**A**) Example of high-power microphotographs of A1 sections stained for PV and mCherry. (**B**) Overall and layer-specific proportion of PV(+)/mCherry(+) cells.

**Supplementary Figure 2.**
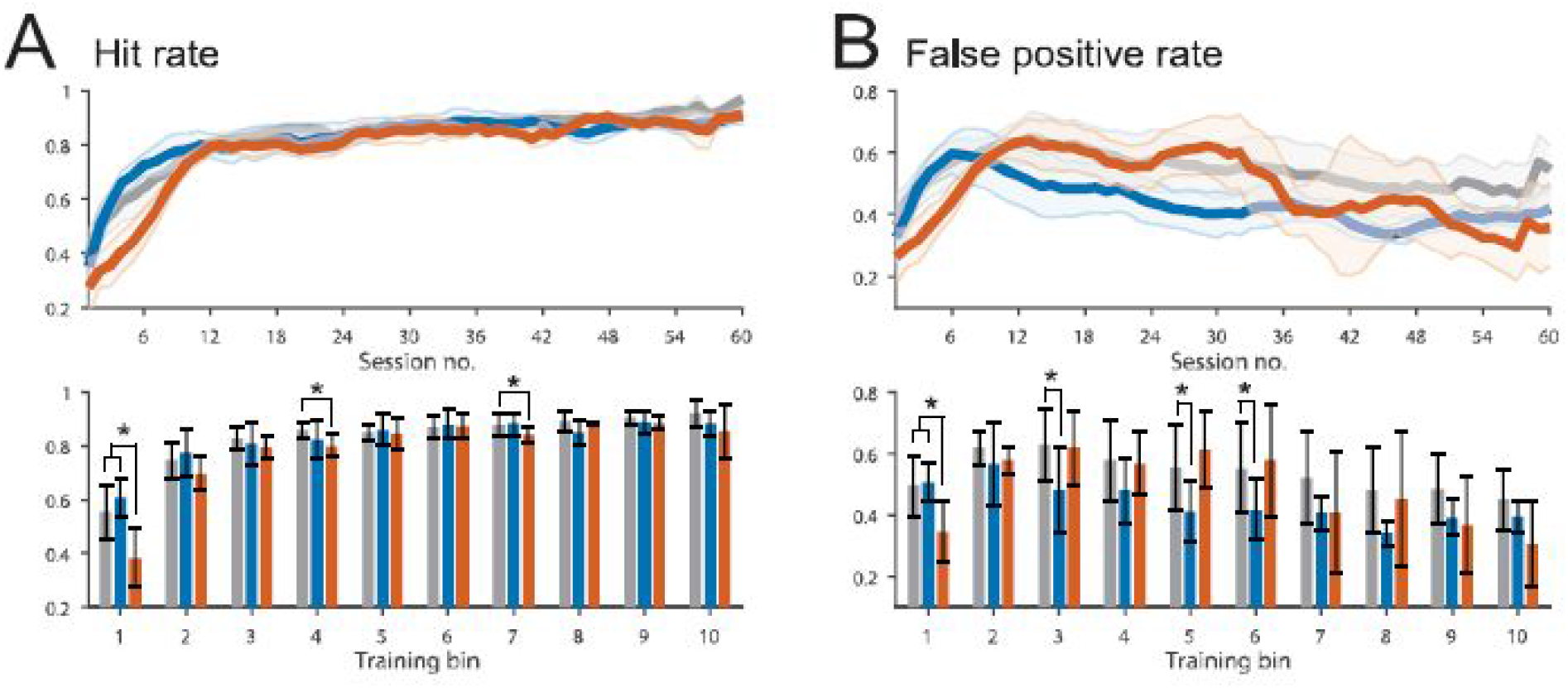
Earlier improvement in false-positive rate accounts for accelerated perceptual learning observed in the PV_I_ group. (**A**) Hit rate and (**B**) false-positive rate for all experimental groups. Main effects of group and group x bin interaction were all significant for both parameters (hit-rate: group, F(2,15) = 29.02, interaction, F(9,18) = 4,69; FP-rate: group, F(2,15) = 5.44, interaction, F(9,18) = 3.01; all p ≤ 0.0167, two-way rmANOVA). FP-rate was lower for the PV_I_ group, relative to the T-Ctrl group for bins 3, 5, and 6 (all p ≤ 0.0367, with Tukey-Kramer test).

**Supplementary Figure 3.**
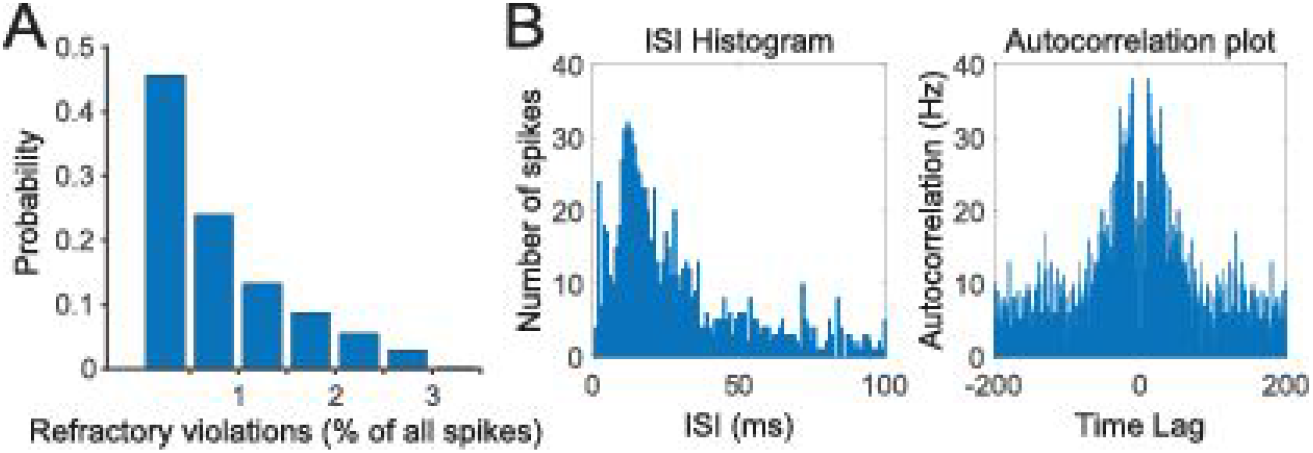
Quality assessment for spike sorting. (**A**) Histogram of the fraction of refractory period violations out of all spikes for identified units for analyses. (**B**) Examples of histograms for an isolated unit of interspike interval (ISI) times (left) and autocorrelation functions (right). Group, number of animals, recording positions, and units: Naïve, 8, 16, 555; T-Ctrl, 7, 14, 491; PV_I_, 6, 12, 531; PV_E_, 8, 16, 507.

**Supplementary Figure 4.**
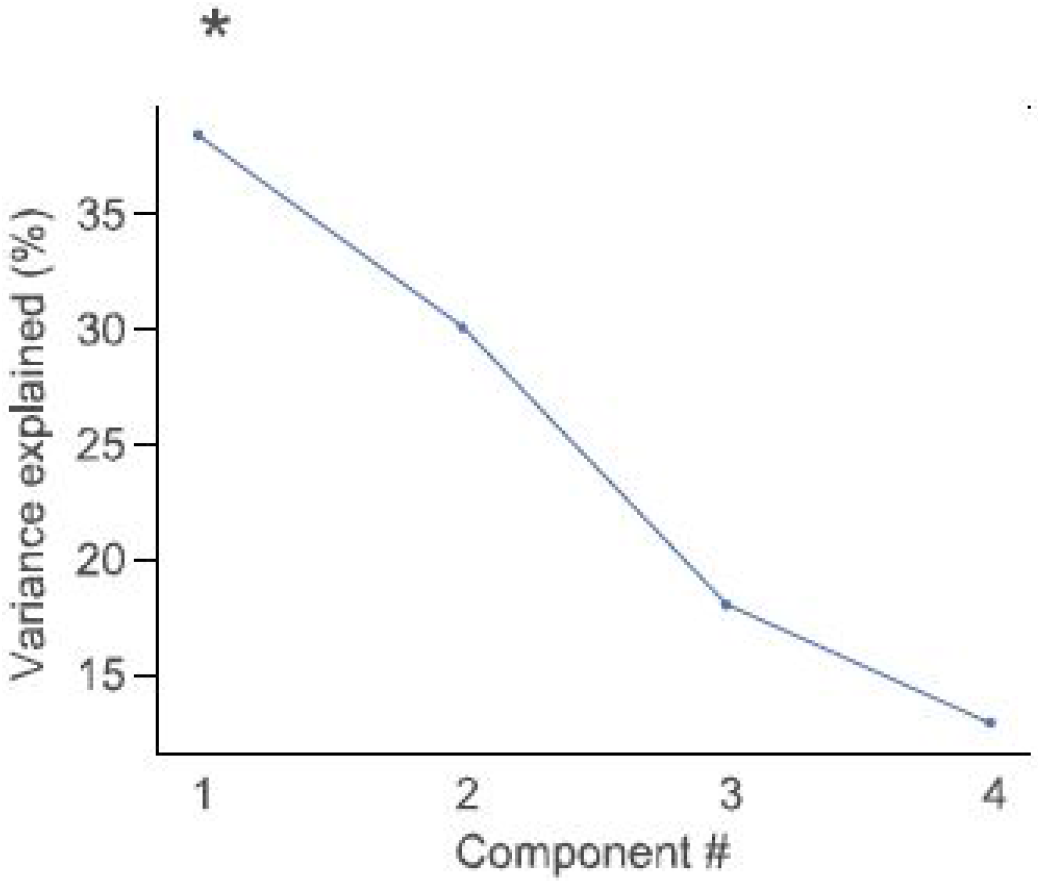
Effect size (variance explained) for each PLS-derived functional component. Permutation sampling was used to perform null-hypothesis significance testing in order to examine the extent to which the variance explained by each component is greater than would be expected by chance. * p ≤ 0.05

